# Optimisation of DNA extraction from nasal lining fluid to assess the nasal microbiome using third-generation sequencing

**DOI:** 10.1101/2024.11.20.624395

**Authors:** Samuel T. Montgomery, Phoebe G. Carr, Jose A. Caparrós-Martín

**Affiliations:** Wal-yan Respiratory Research Centre, Telethon Kids Institute, University of Western Australia, Crawley, Australia; School of Public Health, Curtin University, Bentley, Australia; Curtin Health Innovation Research Institute (CHIRI), Curtin University, 6102, Bentley, Western Australia, Australia; UWA Medical School, The University of Western Australia, 6009, Nedlands, Western Australia, Australia

## Abstract

**Background:** Sampling nasal lining fluid (NLF) via nasosorption is minimally invasive and well tolerated, but the feasibility of assessing the nasal microbiome using this technique is unknown. However, low biomass makes airway samples particularly susceptible to issues related to contaminant DNA. In this study, we evaluated the suitability of DNA isolated using methodologies for low-biomass respiratory samples and assessed how well lining fluid collected via nasosorption captures the nasal microbial diversity and composition compared to the traditional swab sampling approach.

**Methods:** Nasal swabs and NLF were collected from adult volunteers. DNA was extracted from a mock microbial community and NLF using a column-based kit (ZymoBIOMICS), a precipitation-based kit (Qiagen), or a previously published precipitation-based method. Quality and quantity of DNA was assessed and short-read *16S rRNA* sequencing performed to assess feasibility and extraction bias. An optimised extraction methodology was then used to extract DNA from NLF and nasal swabs, and full-length *16S rRNA* sequencing performed to compare microbial profiles between NLF and nasal swabs. Taxonomy was assigned using the nf-core/ampliseq pipeline, the PacificBiosciences/pb-16S-nf pipeline, or the software Emu, and downstream analyses were performed using R packages phyloseq and mixOmics.

**Results:** All extraction methods recovered DNA from the mock community, but only precipitation-based methods yielded sufficient DNA from NLF. Extraction methodologies significantly affected microbial profiles, with mechanical lysis needed to minimize bias against specific genera. Profiles obtained from NLF and swabs were comparable with long-read sequencing.

**Conclusions:** Our findings demonstrate the feasibility of profiling the nasal microbiome using NLF collected via nasosorption and validated two extraction methodologies as suitable for full-length *16S rRNA* sequencing of low-biomass respiratory samples. Our data demonstrate the importance of unbiased DNA extraction methodologies in low-biomass respiratory samples, and the subsequent impact of DNA extraction on observed microbial profiles. Additionally, we demonstrated NLF may be an appropriate surrogate samples for nasal swabs to assess the nasal microbiome using *16S rRNA* sequencing.

## Introduction

The airway microbiota is established early in life and likely plays a major role in host immune responses and mucosal immunology, as evidenced through the functional and compositional microbial variations observed in chronic respiratory diseases (1). The nasal mucosa can be influential in regulating clinical outcomes for respiratory disease – human challenge models of respiratory syncytial virus identified mucosal neutrophil activation as a key factor for development of symptomatic infection (2). The nasal microbiome likely plays a role in regulating the microenvironment, and has been associated with clinical severity of SARS-CoV-2 infection (3). Sampling the nasal microbiome is commonly achieved via nasal or nasopharyngeal swabs, but methods such as nasal lavage allow measurement of proteins in addition to microbial profiling (4). Alternative methods of collecting nasal lining fluid such as nasosorption is well tolerated even in infants, has increased reproducibility and sensitivity for cytokine detection compared to nasal lavage, and avoids the dilution that occurs with nasal lavage (5, 6). However, the feasibility of microbial profiling using nasal lining fluid collected via nasosorption has not been assessed.

A significant barrier for investigating the role of the airway microbiota remains the low bacterial biomass of the airway (7), which makes airway samples particularly susceptible to issues related to contaminant DNA (8, 9). The emergence of accessible and affordable long-read sequencing technologies, particularly Pacific Biosciences (PacBio) and Oxford Nanopore, has opened new frontiers in microbiome research. By amplifying the full length of the *16S rRNA* gene for microbial profiling, taxonomic resolution down to species level and even between strains within a species can be differentiated (10–12), while whole genome long-read sequencing enables the recovery of complete circular bacterial genomes from metagenomic samples (13, 14). As a result, extracting high quality, high molecular weight DNA from metagenomic samples while maintaining high efficiency for low biomass respiratory samples is vital. Common methods for genomic DNA extraction and sequencing for microbial profiling have been developed, optimised, and benchmarked for use with high microbial biomass such as faecal samples (15, 16). While there are published methods for efficient extraction of DNA from samples such as lung tissue or bronchoalveolar lavage fluid for *16S rRNA* sequencing or metagenomic sequencing (17–21), these methodologies have not been evaluated for long-read sequencing that requires high molecular weight DNA. In this study, we evaluated the suitability of DNA isolated using methodologies for low-biomass respiratory samples and assessed how well lining fluid collected via nasosorption captures the nasal microbial diversity and composition compared to the traditional swab sampling approach.

## Methods

Full details are available in the Supplementary Information.

### Study population

Nasal samples were obtained from adult volunteers through a human research ethics committee approved study (CAHS HREC #RGS1470). This study included samples from 19 adults (male 78%). All participants provided written informed consent.

### Nasal sampling

Nasal lining fluid and nasal swabs were used in this study. Nasal lining fluid was obtained via nasosorption using a nasosorption device with a synthetic absorptive matrix (SAM) (Mucosal Diagnostics, UK) as previously described (22). Briefly, the device was inserted into the nasal lumen and contact made with the inferior turbinate for 60 seconds before removal into the associated cryotube and stored at −80°C. Nasal swabbing was performed by swabbing the nasal inferior turbinate with a flexible shaft flocked swab (Copan Diagnostics, USA) collected in Puregene Lysis Buffer (Qiagen, USA).

### Elution of nasal lining fluid

Nasal lining fluid was eluted from nasosorption devices as previously described (22) with slight modifications dx.doi.org/10.17504/protocols.io.14egn3nn6l5d/v1). Briefly, nasosorption devices were thawed on ice and the SAM was submerged in 300μL of sterile phosphate buffered saline (ThermoFisher, USA) containing 0.05% (v/v) Tween20 (Sigma, USA) and vortexed for 30 seconds. The moist SAM was then added to a spin filter column (SpinX, Corning, USA) and centrifuged at 16,000g for 30 minutes at 4°C. The eluate was then split, with 255μL of the supernatant removed and stored for protein analysis, and the remaining 45μL and pellet frozen at −80°C for DNA extraction.

### DNA extraction

For optimisation, DNA was extracted from nasal lining fluid (NLF) and a mock community of bacteria (Zymo Research, USA) using the ZymoBIOMICS DNA Mini kit (ZYMO), the Qiagen Puregene Blood kit (QIA), or a previously published methodology utilising choroform:isoamyl alcohol and polyethylene glycol (PEG) (19). Extraction methods utilised either mechanical lysis via bead beating with (MP) and without (BB) enzymatic lysis. Bead beating utilised the FastPrep-24 (MP Bio) or the Precellys 24 (Bertin Technologies, USA) tissue homogenisers, with 0.1mm and 0.5mm glass beads from the ZymoBIOMICS DNA Mini kit. Enzymatic lysis utilised MetaPolyzyme (Sigma, USA), a mix of six lytic enzymes developed by the Extreme Microbiome Project (23). For comparisons between NLF and nasal swabs, DNA was extracted using an optimised methodology (dx.doi.org/10.17504/protocols.io.3byl4qwqovo5/v1). DNA was quantified using a Nanodrop spectrophotometer and a Qubit fluorometer, and fragment sizes assessed using a TapeStation Genomic DNA ScreenTape (Agilent, USA).

### Illumina 16s rRNA sequencing

For short-read *16S rRNA* amplicon-based microbial profiling, extracted DNA was amplified and sequenced by the Australian Genome Research Facility. Briefly, forward (341F: *CCTAYGGGRBGCASCAG*) and reverse (806R: *GGACTACNNGGGTATCTAAT*) primers were used to amplify the V3-V4 hypervariable region of the *16S rRNA* gene, amplicons barcoded and combined using equimolar pooling, and sequenced (2×300bp paired-end) using the Illumina MiSeq. Raw data was processed using the nf-core/ampliseq pipeline (24, 25) which assigns taxonomy to amplicon sequence variants (ASVs) using dada2 (26) with the SILVA (v138) database.

### Oxford Nanopore 16S rRNA sequencing

For full-length *16S rRNA* sequencing, extracted DNA was amplified using the NanoID kit (Shoreline Biome, USA) targeting a 2.5kB fragment of 16S rRNA-ITS-23S RNA region. Library preparation was performed using a ligation sequencing kit (SQK-LSK114, Oxford Nanopore) and sequenced using a R.10.4.1 flow cell on the MinION Mk1C (Oxford Nanopore). Raw data were basecalled in super-accurate mode using Dorado v0.4.0 (https://github.com/nanoporetech/dorado) with the dna_r10.4.1_e8.2_400bps_sup@v4.2.0 model. Reads were filtered for length (>2000bp & <3000bp) and quality (>Q10) using chopper v0.5.0 (https://github.com/wdecoster/chopper), and read statistics visualised using NanoPlot v1.33.0 (https://github.com/wdecoster/NanoPlot). Taxonomy was assigned to the filtered reads using Emu v3.4.5 (12) with the default database (rrnDB v5.6 & RefSeq).

### PacBio *16S rRNA* sequencing

For full-length *16S rRNA* sequencing, extracted DNA was amplified and sequenced by the Australian Genome Research Facility. Briefly, forward (F27: *5’GCATC/barcode/AGRGTTYGATYMTGGCTCAG3’*) and reverse (R1492: *5’GCATC/barcode/RGYTACCTTGTTACGACTT3’*) primers were used to amplify the full (V1-V9) *16S rRNA* gene and sequenced using a PacBio Revio. Raw data was processed using the PacificBiosciences/pb-16S-nf pipeline (https://github.com/PacificBiosciences/pb-16S-nf) which assigns taxonomy to amplicon sequence variants (ASVs) using dada2 with the GTDB (r207) and SILVA (v138) databases.

### 16S rRNA sequencing data analysis

Contaminating sequences were removed using the decontam R package (6), alpha diversity assessed using the phyloseq R package and abundance counts were centre log ratio transformed for further analysis using the MixMC framework implemented in the mixOmics R package, with principal component analysis (PCA) and sparse PLS discriminant analysis (sPLS-DA) used to identify a signature of discriminative ASVs associated with DNA extraction methods (7, 8).

### Statistical analysis

All statistical analysis was performed using the R statistical language (v4.1.2) (27). For data wrangling and visualisation we used the tidyverse suite of R packages (28), specifically the ggplot2 R package (29), or GraphPad Prism (v9.1; GraphPad Software, USA). Principal component analysis plots were generated using the mixOmics R package (30).

### Data availability

All data required to reproduce the results of this study are available within the manuscript and supplementary information. Demultiplexed sequencing data are deposited in the NCBI Sequencing Read Archive and available under the BioProject PRJNA1185436. All code used to analyse the data is available online: https://github.com/samuelmontgomery/nlf_optimisation

## Results

### Precipitation-based DNA extraction methods have higher yield and higher DNA quality than the silica-column based method

All extraction methods successfully extracted DNA from the mock community (Figure 1A). Both the ZYMO and QIA kits resulted in a significantly lower amount of DNA recovered when compared with the PEG method. Both ZYMO and QIA resulted in a 260/280 ratio significantly different from the optimal ratio of 1.8 (Figure S1A), and all methods resulted in a 260/230 ratio significantly lower than the optimal ratio of 2.0 (Figure S1B). When assessed for fragment size, DNA extracted using QIA had a significantly higher peak molecular weight compared to ZYMO and PEG (p = 0.0043 and p = 0.0024 respectively; Figure 1B). All extraction methods successfully extracted DNA from NLF (Figure 1C), however QIA extracted significantly more DNA than ZYMO (p < 0.05). Both ZYMO and QIA resulted in a 260/280 ratio significantly different from the optimal ratio of 1.8 (Figure S1C), and all methods resulted in a 260/230 ratio significantly lower than the optimal ratio of 2.0 (Figure S1D). When assessed for fragment size, DNA extracted from NLF using ZYMO was too low in concentration to successfully quantify fragment sizes, and there was no significant difference in molecular weight of extracted DNA between QIA and PEG (Figure 1D).

**Figure 1:**
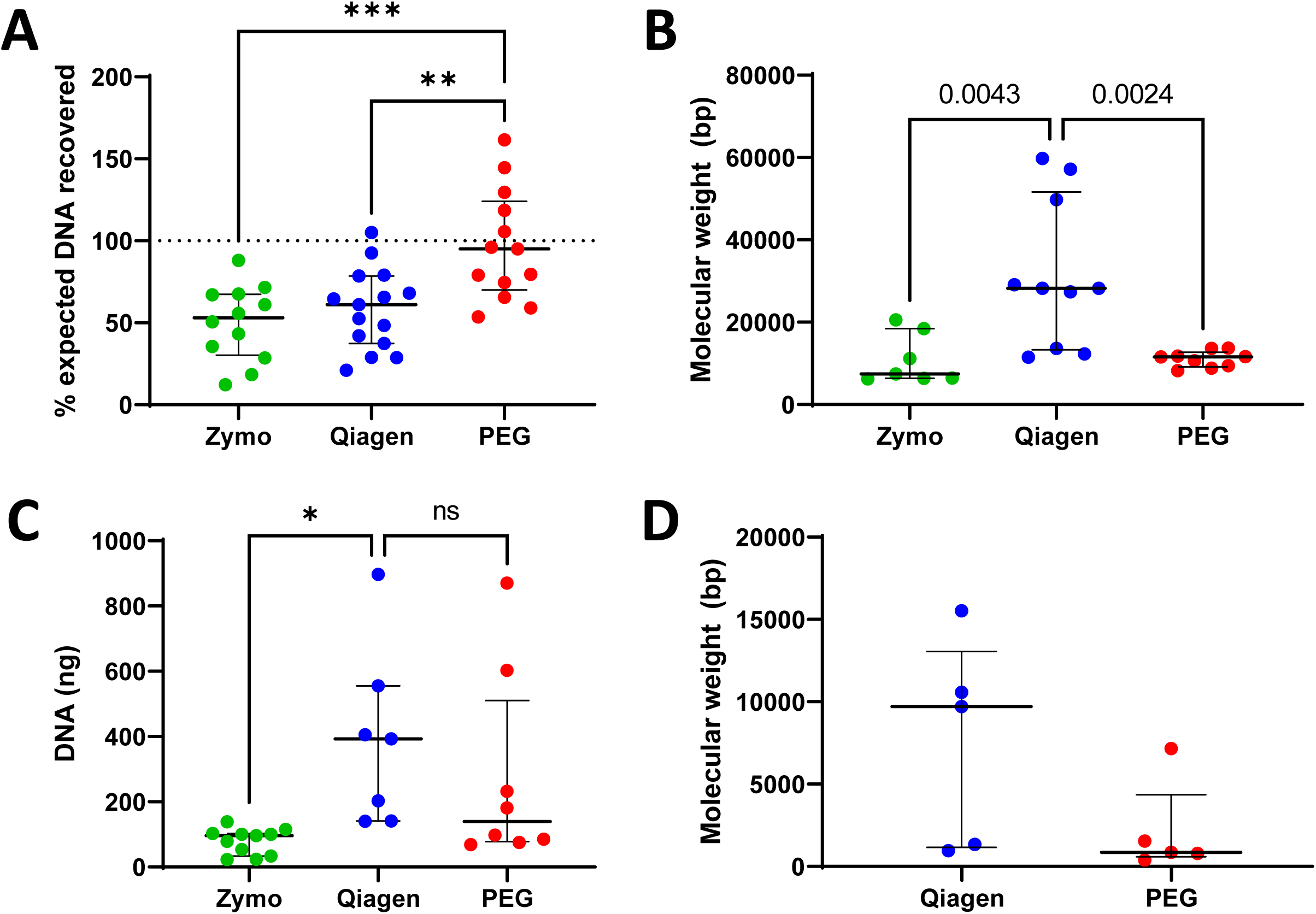
Quantity and size of DNA extracted using Zymo, Qiagen, and PEG methods. **(A)** Percentage of expected DNA yield from the mock community standard. **(B)** Peak molecular weight of DNA extracted from the mock community standard. **(C)** Quantity of DNA extraction from nasal lining fluid. **(D)** Peak molecular weight of DNA extracted from nasal lining fluid. * p<0.05; ** p<0.01; *** p<0.001; **** p<0.0001. Dotted line represents expected values. Differences between groups assessed using one-way ANOVA with Tukey’s test for multiple comparisons.

### Extracted DNA could be amplified for *16S rRNA* sequencing

To assess the suitability of extracted DNA for microbial profiling, the V3-V4 region of the *16S rRNA* gene was amplified. Amplicon libraries from 40 samples, including 5 negative extraction controls, were generated and sequenced at the Australian Genome Research Facility (Melbourne, Australia). After sequencing, 1,911,296 paired-end reads were obtained with high base calling accuracy, with a median Phred quality score of 36.10 (IQR: 32.98– 37.03). After filtering data to remove singlets, there were 305 ASVs representing bacterial taxa, of which 107 ASVs were present in negative extraction and sequencing controls. Negative extraction controls were clearly separated from true samples by library size (Figure S2A) and clustered separately from true samples (Figure S2B). Data was then filtered to remove taxa with less than 5 reads in at least two samples and potential contaminants filtered using the prevalence method from the decontam R package (31), resulting in 70 ASVs remaining in the cleaned dataset. When the taxa in the unfiltered dataset were compared to the filtered dataset using principal coordinate analysis, there was no major change in overall structure of the data as confirmed by Procrustes analysis (sum of squares = 0.00009053; correlation = 1.00; p = 0.001).

### Mechanical lysis reduced extraction bias

To assess potential bias in DNA extraction, a mock community of known composition was extracted in parallel with all three extraction methods, using either enzymatic lysis alone or in combination with mechanical lysis and submitted for *16S rRNA* sequencing. Extraction with enzymatic lysis alone, independent of extraction method, resulted in a loss of gram-positive genera *Staphylococcus*, *Limosilactobacillus*, and *Listeria*, and resulted in an overrepresentation of gram-negative genera *Bacillus* and *Escherichia-Shigella* (Figure 2A). When alpha diversity measures (Shannon, Simpson) were assessed, DNA extracted using enzymatic lysis alone clearly resulted in lower diversity (Figure 2B). As expected, ZYMO with mechanical lysis resulted in a diversity closely matching the profile of the reference DNA, while PEG and QIA with mechanical lysis were not significantly different from control alpha diversity measures (Figure 2B). Multivariate analysis using sPLS-DA (Figure 3A) identified that samples extracted with enzymatic lysis alone were associated with all four gram-negative genera present in the mock community, while mechanical lysis was associated with all four gram-positive genera (Figure 3B). However, while samples cluster by extraction kit, there was overlap between all three methods with mechanical lysis (Figure 3A). When only the mechanical lysis samples were included in analysis, samples clustered by which bead beater was used for lysis rather than by extraction kit (Figure 3C). The sPLS-DA analysis identified the gram-negative genus *Bacillus* associated with the Precellys, while gram-positive genus *Limosilactobacillus* was associated with the FastPrep-24 (Figure 3D).

**Figure 2:**
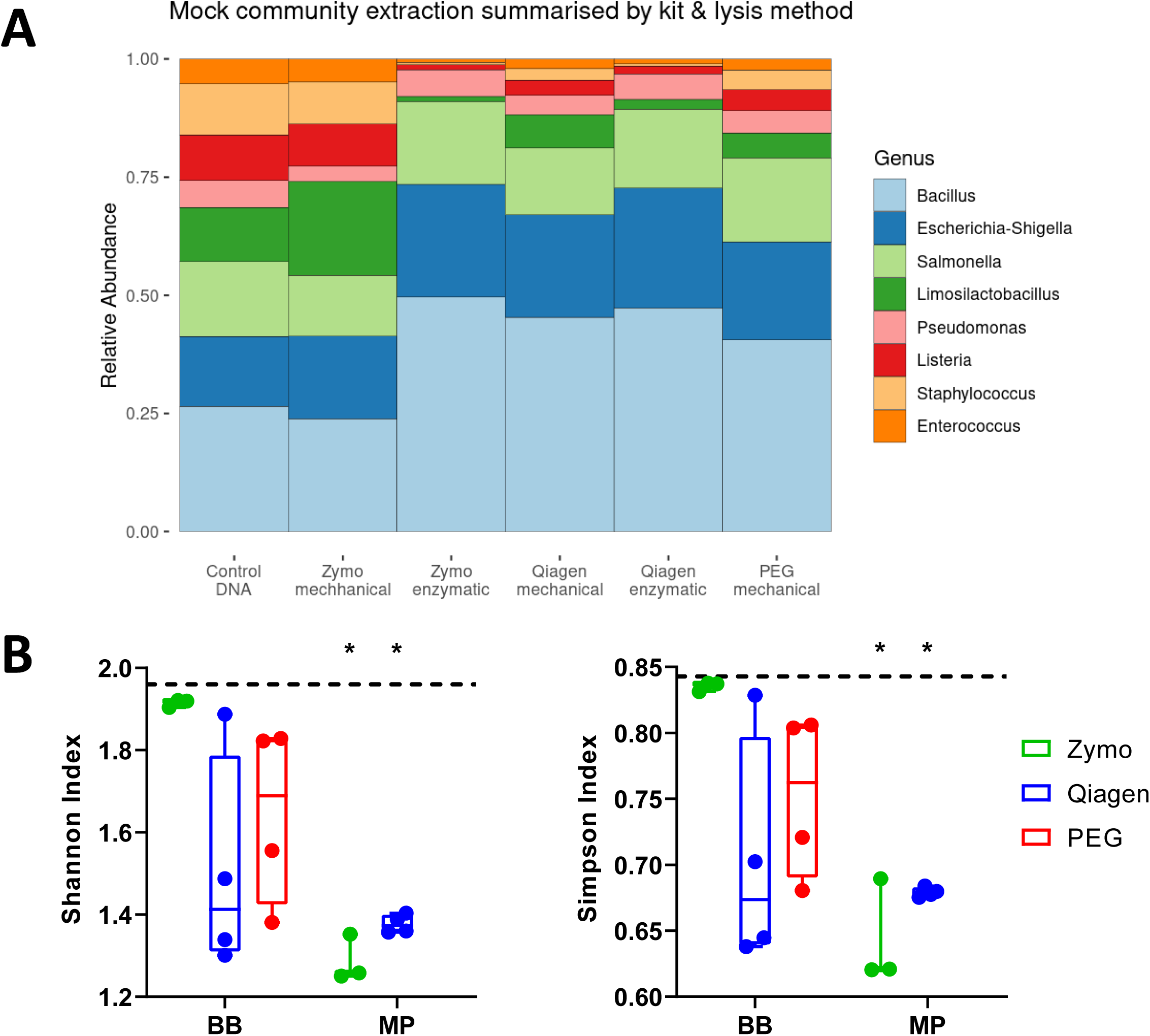
Relative abundance and alpha diversity of DNA extracted from a mock community standard by extraction and lysis method. **(A)** The relative abundance of the 8 taxa present in the mock community, separated by extraction kit and lysis method. **(B)** Alpha diversity metrics (Shannon, Simpson) of mock community standard samples by extraction kit and lysis method. * p<0.05. Dotted line represents values obtained for control DNA. Comparisons to control DNA diversity used a one-way t-test for statistical significance.

**Figure 3:**
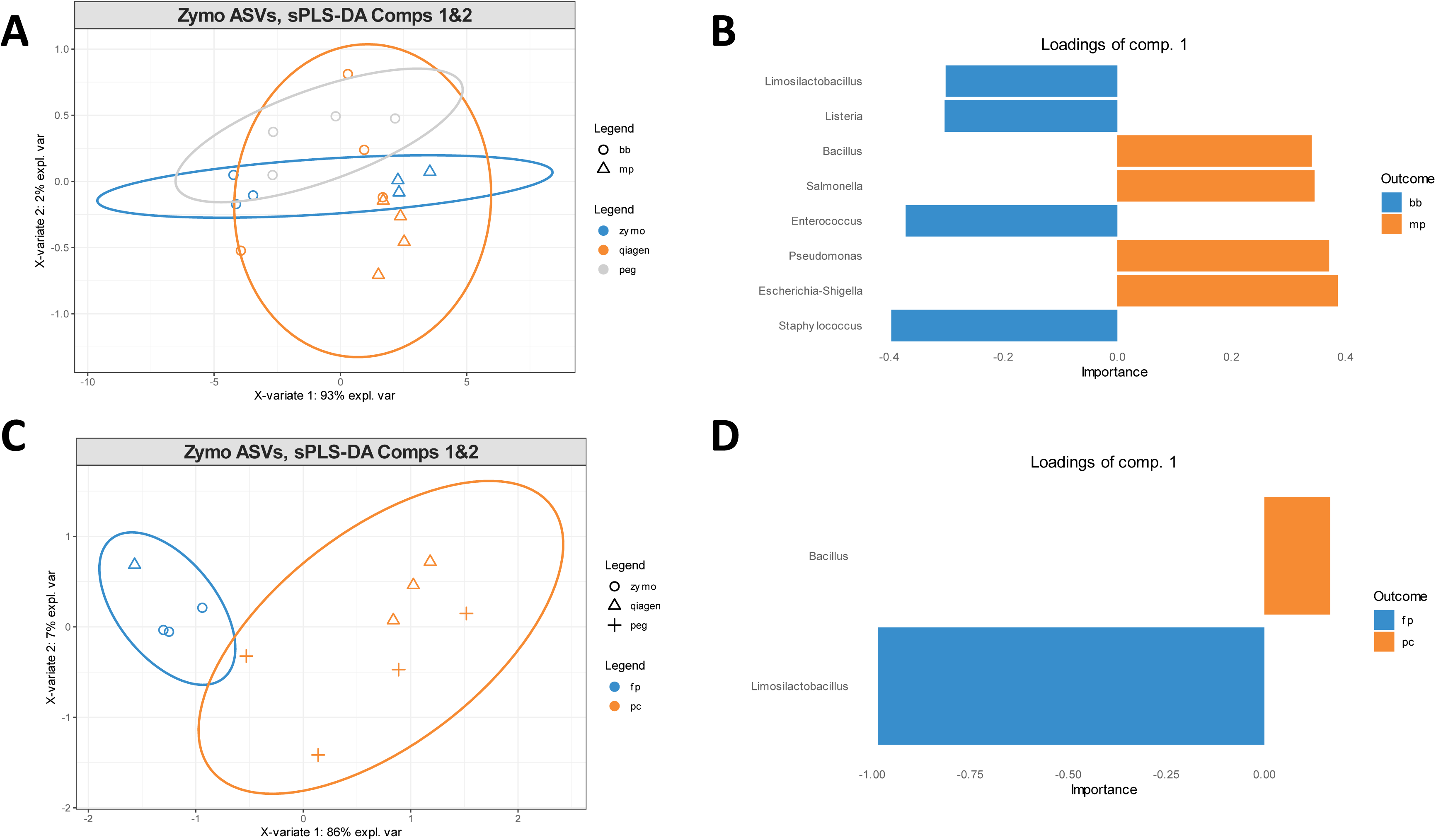
Identification of a bacterial signature in a mock community discriminating lysis method in during DNA extraction with sPLS-DA. **(A)** sPLS-DA sample plot of ASVs from mock community samples on the first two components separated by extraction (colours) and lysis (shape) methodologies with 95% confidence level ellipse plots **(B)** Loading plot of top contributing genera to the first component of the sPLS-DA analysis in (A). **(C)** sPLS-DA sample plot of ASVs from mock community samples on the first two components separated by bead beating device (colours) and extraction (shape) methodologies with 95% confidence level ellipse plots **(D)** Loading plot of top contributing genera to the first component of the sPLS-DA analysis in (C).

### Nasal microbial profiles can be obtained from nasal lining fluid

To assess the feasibility of profiling the nasal microbiome using nasosorption, NLF was pooled and extracted in parallel with all three extraction methods. Due to the low quality and quantity of DNA extracted from nasal lining fluid with ZYMO, only DNA extracted with QIA and PEG was submitted for *16S rRNA* sequencing. Microbial profiles from NLF using both extraction methods were dominated by *Corynebacterium*, with a low abundance of other canonical airway microbes (Figure 4A). When alpha diversity measures (Shannon, Simpson) were assessed, DNA extracted using PEG resulted in slightly higher diversity represented in the samples, but diversity was low overall in NLF (Figure 4B). Multivariate analysis using sPLS-DA resulted in samples extracted via PEG and QIA clustering separately (Figure 5A) and identified a signature of genera, including *Haemophilus, Dolosigranulum,,* and *Lawsonella* associated with PEG, while samples extracted using QIA were associated with genera including *Prevotella, Klebsiella, Moraxella, Streptococcus* (Figure 5B-C). PERMANOVA analysis showed significant differences between PEG and QIA (pseudo-F = 7.7596, *p* = 0.002, R^2^ = 0.37378), indicating that the extraction methodology significantly affects the overall composition of the microbial communities in nasal lining fluid, however as both sample types were dominated by *Corynebacterium*, this likely represents small differences in low abundant taxa observed.

**Figure 4:**
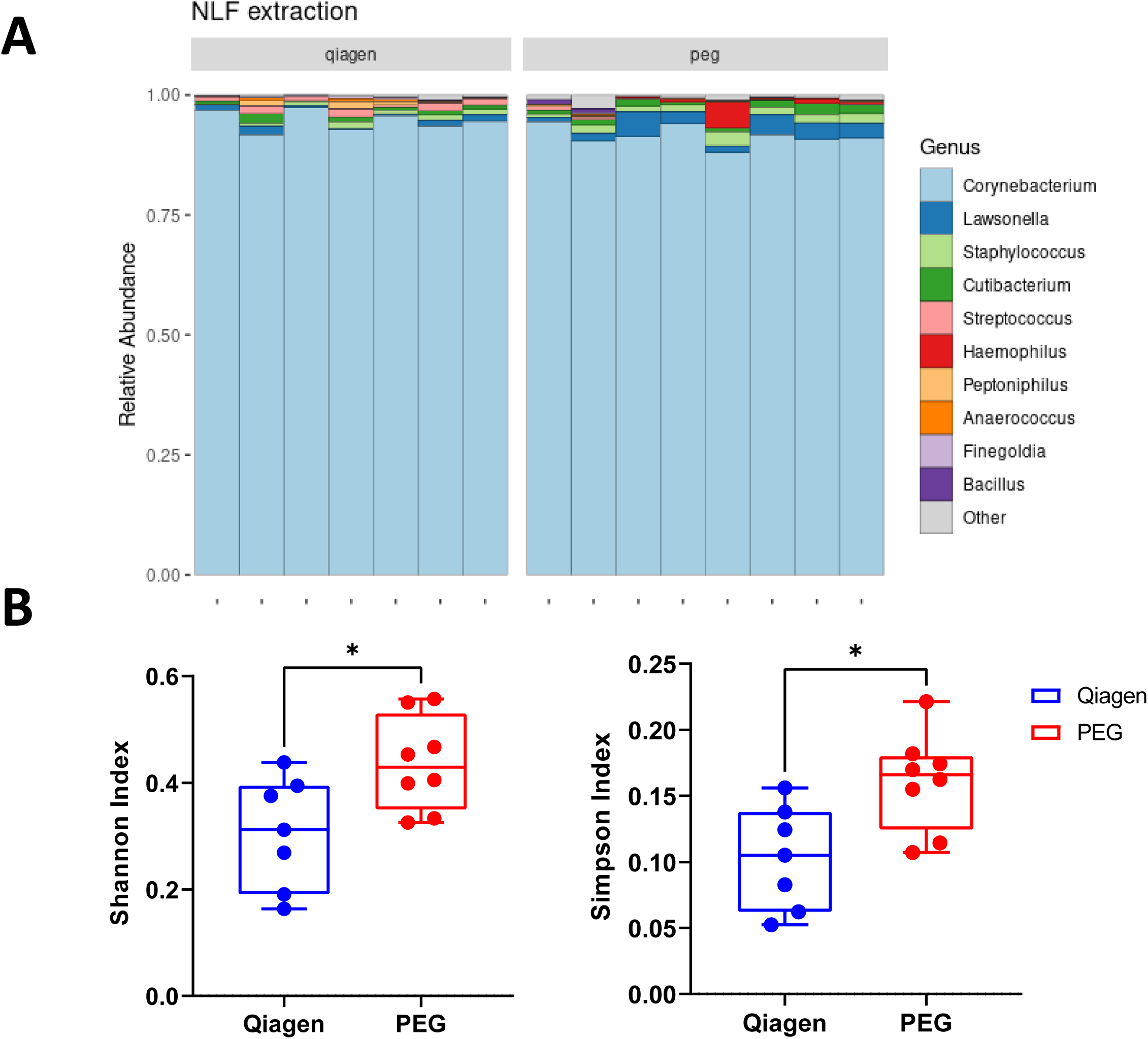
Relative abundance and alpha diversity of DNA extracted from nasal lining fluid samples. **(A)** Relative abundance of nasal lining fluid samples separated by extraction kit. **(B)** Alpha diversity metrics (Shannon, Simpson) of nasal lining fluid samples separated by extraction kit (red: PEG, blue: Qiagen). * p<0.05. Comparisons between extraction kits used a Mann-Whitney U-test for statistical significance.

**Figure 5:**
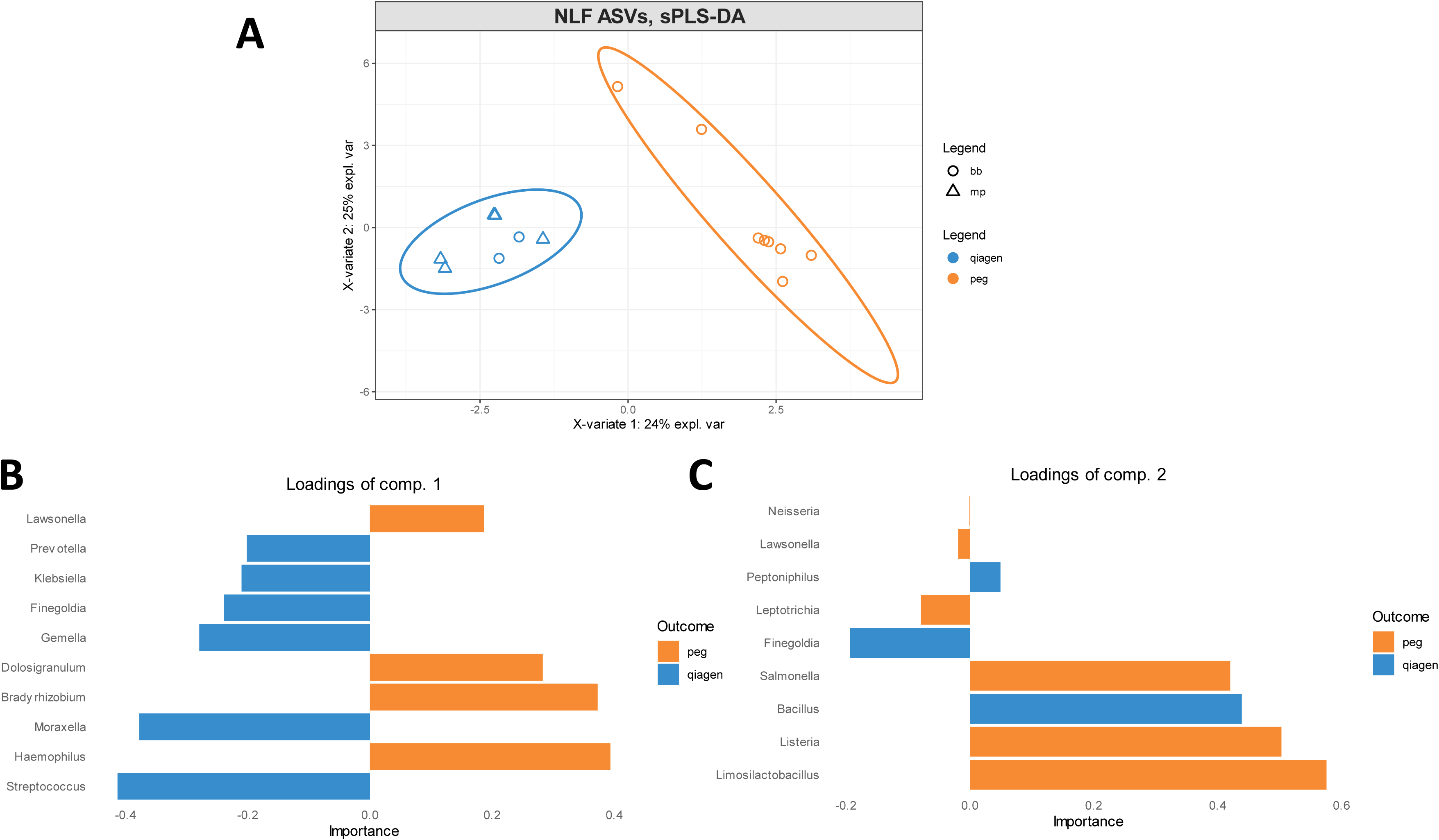
Identification of a bacterial signature discriminating extraction kit during DNA extraction from nasal lining fluid with sPLS-DA. **(A)** sPLS-DA sample plot of ASVs from nasal lining fluid samples on the first two components separated by extraction kit (colours) and lysis method (shape) with 95% confidence level ellipse plots. **(B)** Loading plots of top contributing genera to the first and **(C)** second components of the sPLS-DA analysis

### Microbial profiles from nasal lining fluid are representative of profiles obtained from nasal swabs

As nasal swabbing is a well-established technique for assessing the nasal microbiome, paired nasal swabs and nasal lining fluid obtained from separate nostrils on the same participant were assessed to determine whether nasal lining fluid is an appropriate surrogate sample. Microbial profiles were obtained using full-length *16S rRNA* sequencing across two long-read sequencing technologies PacBio Revio (PB) and Oxford Nanopore using a Shoreline Biome NanoID library (SL). Both SL (Figure S4) and PB (Figure S5) were able to assign taxonomy to species-level resolution, with similar microbial profiles observed in both NLF and swabs (Figure 6). Samples sequenced with PB had less *Corynebacterium* observed compared to SL, which may be due to biases introduced during amplification as SL targets the ITS and part of *23S rRNA* gene in addition to *16S rRNA* gene. When projected into two-dimensional space via PCA, a significant overlap between NLF and nasal swabs were observed when SL and PB are combined (Figure 7A) or analysed separately (Figures 7B-C), however SL and PB clustered separately from each other. PERMANOVA analysis found no significant differences between microbial community composition obtained from NLF and swabs (pseudo-F = 0.3987, p = 0.992, R^2^ = 0.03834). Multivariate analysis using sPLS-DA (Figure 8) identified multiple bacterial species associated with nasal swabs in both PB and SL data, such as *Streptococcus sanguinis* and *Staphylococcus pasteuri* (Figure 8B & 8D). When alpha diversity measures (Shannon, Simpson) were assessed, both sample types resulted in similar diversity with minor differences observed between SL and PB data (Figure S8).

**Figure 6:**
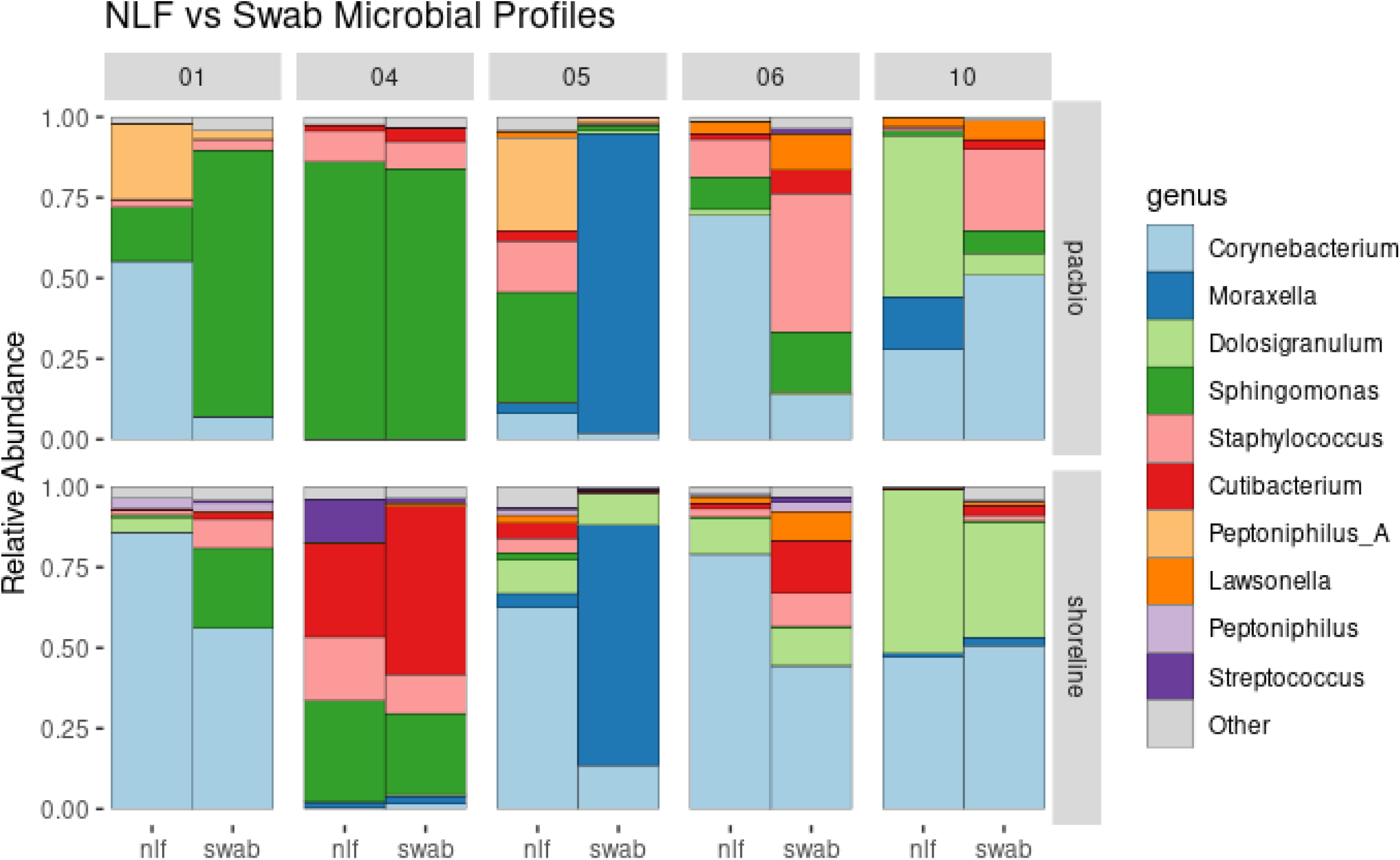
Effects of sample collection and library preparation on composition of the nasal microbiome. Relative abundance of taxa present in paired nasal lining fluid and nasal swab samples, with PacBio data on the top row and Shoreline data on the bottom row, and participant ID above each pair of columns.

**Figure 7:**
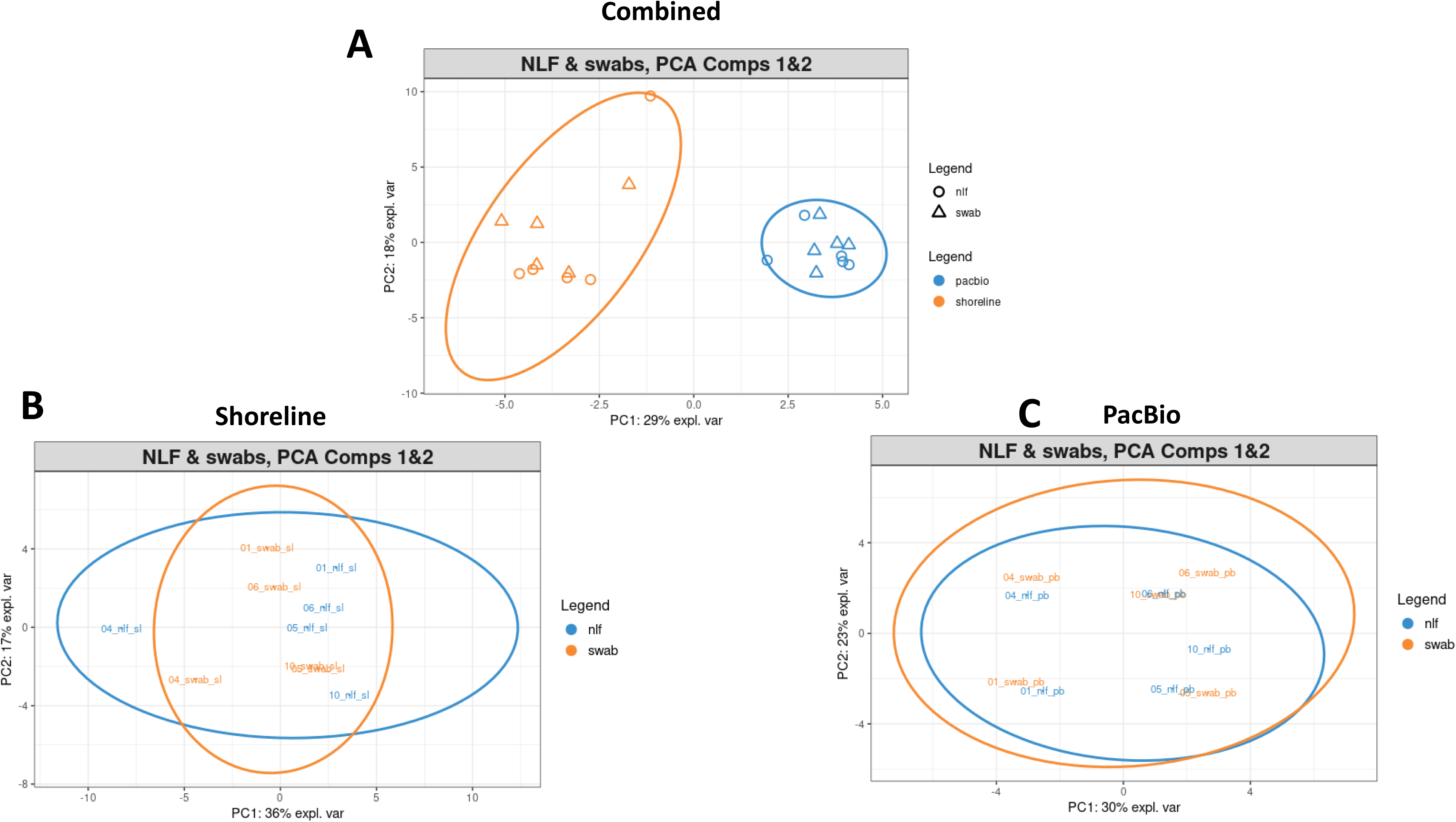
Principal component analysis of taxa from paired nasal swabs and nasal lining fluid samples. **(A)** When analysed together, taxa present in nasal swab and nasal lining fluid samples clustered by library preparation, and not collection method. When analysed separately, there was significant overlap between taxa present in nasal swab and nasal lining fluid samples sequenced using **(B)** Shoreline and **(C)** PacBio libraries. Ellipses represent 95% confidence intervals.

**Figure 8:**
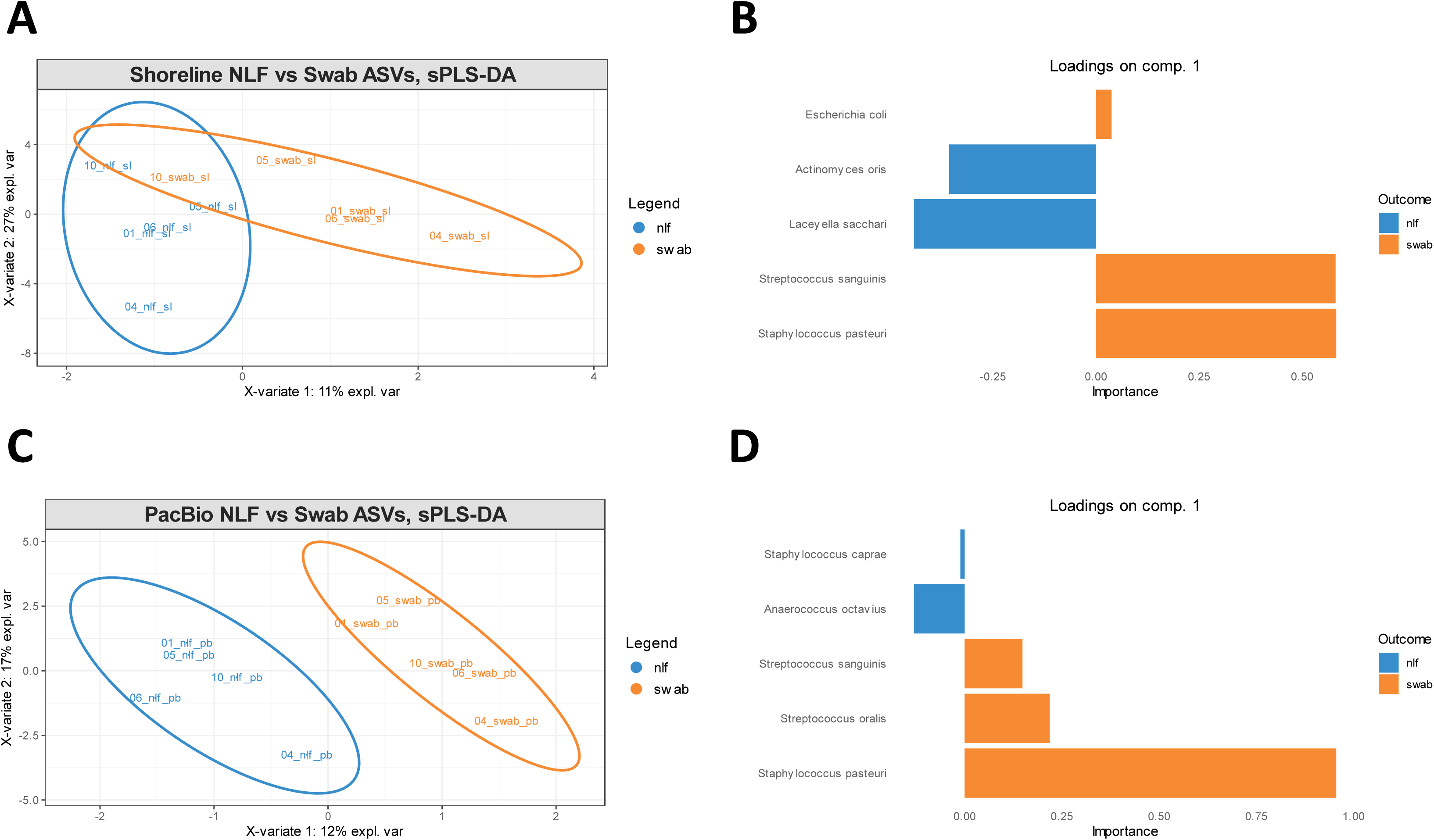
Multivariate analysis of paired nasal swabs and nasal lining fluid identify a microbial signature associated with sampling method with sPLS-DA. **(A)** sPLS-DA sample plot of taxa identified by Shoreline library preparation in nasal swab and nasal lining fluid samples on the first two components with 95% confidence interval ellipse plots. **(B)** Loading plots of top contributing taxa to the first component of the sPLS-DA analysis in (A). **(C)** sPLS-DA sample plot of taxa identified by PacBio library preparation in nasal swab and nasal lining fluid samples on the first two components with 95% confidence interval ellipse plots. **(D)** Loading plots of top contributing taxa to the first component of the sPLS-DA analysis in (C).

## Discussion

In the present study, we demonstrate the feasibility of profiling the nasal microbiome using nasal lining fluid collected via nasosorption and validated two extraction methodologies as suitable for full-length *16S rRNA* sequencing of low-biomass respiratory samples. Our data demonstrate the importance of unbiased DNA extraction methodologies in low-biomass respiratory samples, and the subsequent impact of DNA extraction on observed microbial profiles. Additionally, we demonstrated nasal lining fluid may be an appropriate surrogate samples for nasal swabs to assess the nasal microbiome using *16s rRNA* sequencing.

All three extraction methodologies assessed in this study were able to extract DNA from the mock community of known composition, however only the precipitation-based methods were able to recover a sufficient quantity of DNA from nasal lining fluid. All extraction methods resulted in spectroscopic measures for DNA quality (260/280 and 260/230 ratios) that deviate from the recommended values for optimal downstream usage but had no effect on the feasibility of conducting *16S rRNA* sequencing on the samples. We found significant differences in microbiome composition between extraction kits/methods using a mock community of known composition, with mechanical lysis required to reduce bias against harder-to-lyse genera. Similarly, the methodology/kit for DNA extraction significantly influenced the microbial profiles obtained from nasal lining fluid. Prior studies have shown variability in microbiome data can heavily depend on the DNA extraction and lysis methodology (32–34), and our results corroborate these findings in both a mock community of bacteria and paired low-biomass respiratory samples, targeting both the V3-V4 hypervariable regions of the *16S rRNA* gene, and the full-length *16S rRNA* gene using third generation long-read sequencing.

We compared the microbial profiles obtained from nasal lining fluid to nasal swabs, the gold standard sample for assessing nasal microbiome, using third generation long-read sequencing enabling amplification of the entire *16S rRNA* gene. Microbial profiles have previously been compared between nasal lining fluid and nasal lavage (35), with similar genera identified in nasal lining fluid as this study, but with no positive control or mock community to confirm unbiased DNA extraction. Similarly, microbial profiles obtained from nasal lavage has been compared to nasal swabs in children with cystic fibrosis (4), and found nasal swabs were more likely to detect *Staphylococcus* species – which was also observed in this study. This may be due to the differences in sampling methods, where nasosorption relies on bacteria to be present in the lining fluid, nasal swabs scrape the epithelial lining of the airway and may sample more bacteria that rely on attaching to host surfaces.

This study had several limitations. Firstly, we did not include a magnetic bead-based DNA extraction protocol which are becoming more commonly used in the literature and cannot rule out that they may perform similarly in extraction efficiency. Secondly, as we only performed full-length *16S rRNA* sequencing and did not quantify the bacterial load in the respiratory samples, we cannot determine differences in recovery of host DNA between protocols, or how differences in biomass between samples may have impacted the microbial profiles observed. Future work will include whole metagenomic sequencing to assess differences observed without amplification bias and quantify any differences in non-bacterial members of the nasal microenvironment (e.g., fungi, archaea).

In summary, our findings demonstrate a methodology for DNA extraction from low-biomass respiratory samples such as nasal lining fluid collected via nasosorption, and suitability as a surrogate sample for nasal swabs to assess the nasal microbiome via *16S rRNA* sequencing. Future use of this methodology will facilitate more frequent sampling of the nasal microenvironment while potentially reducing the invasiveness for research study participants.

---

**Supplementary Fig 1.**
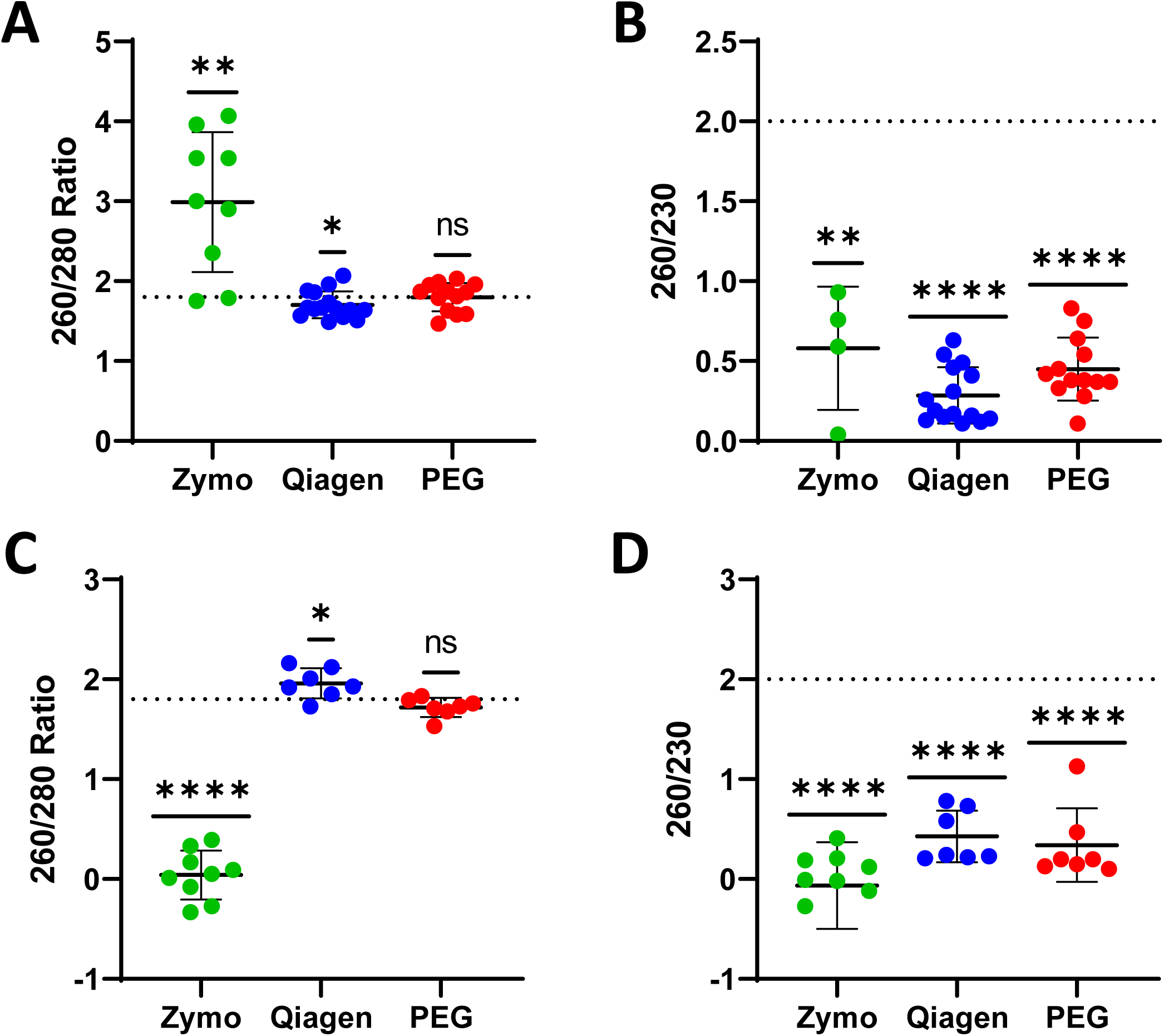

**Supplementary Fig 2.**
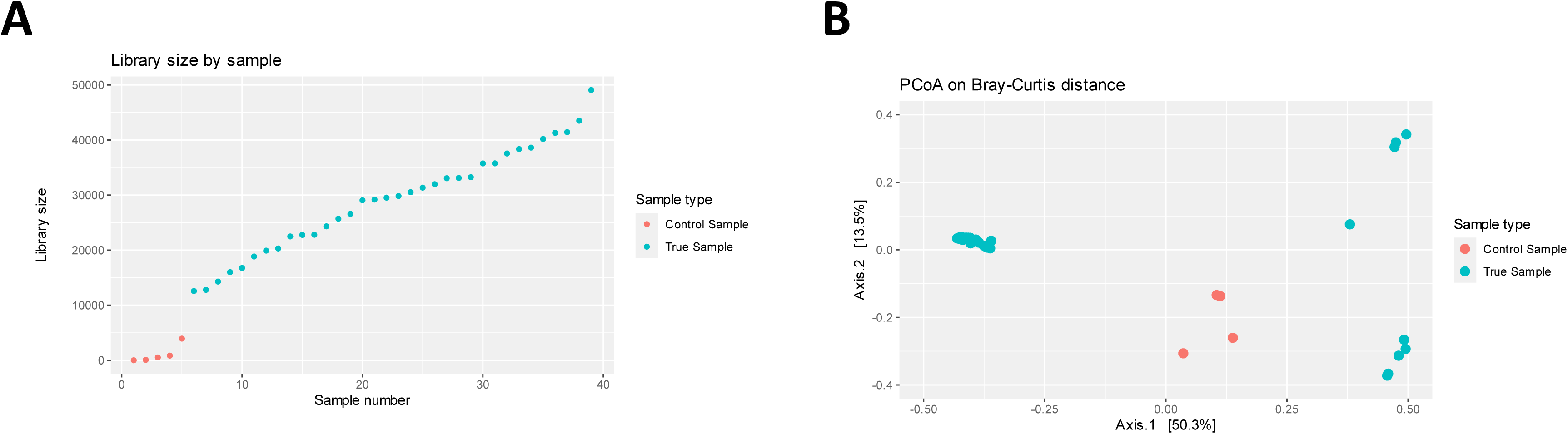

**Supplementary Fig 3.**
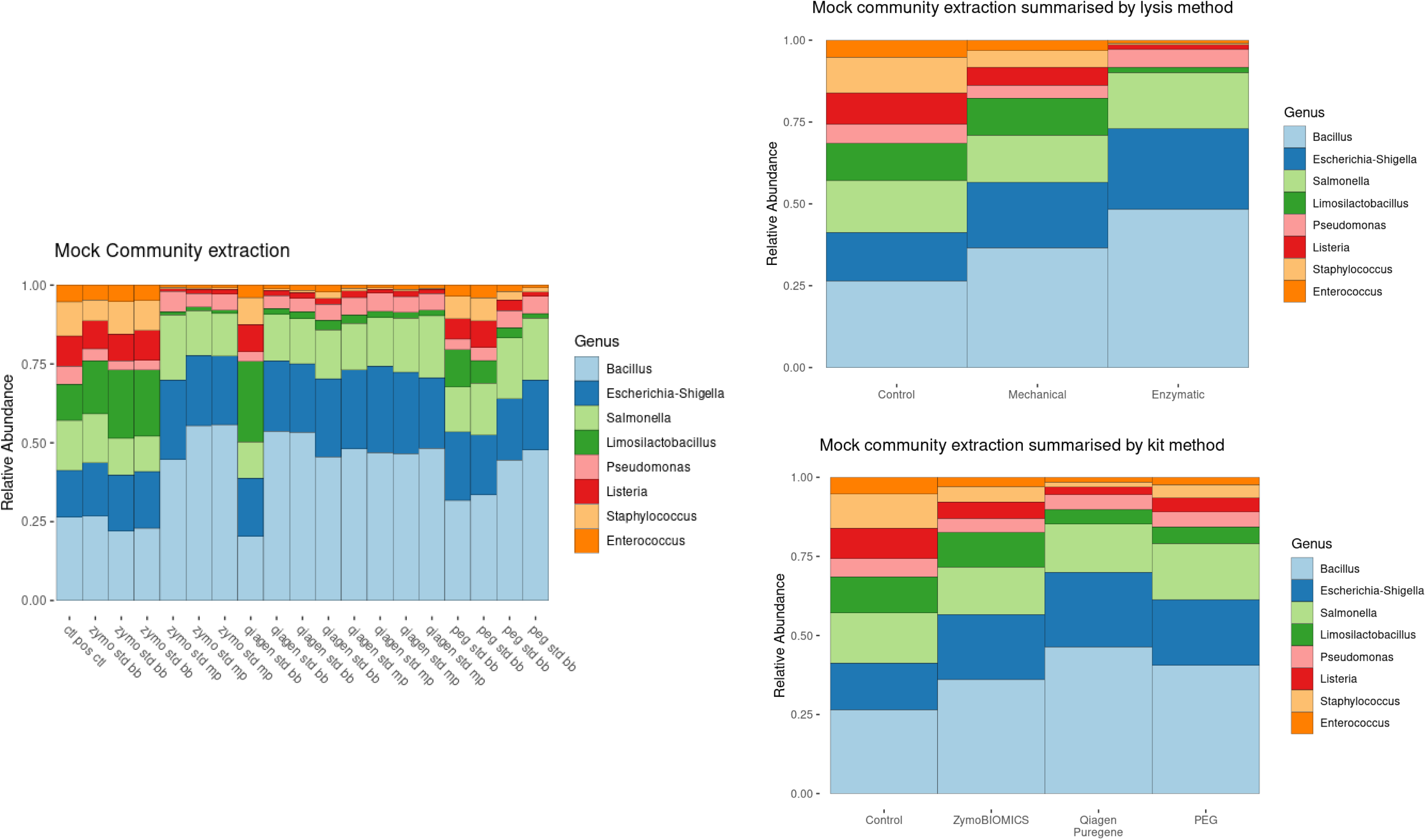

**Supplementary Fig 4.**
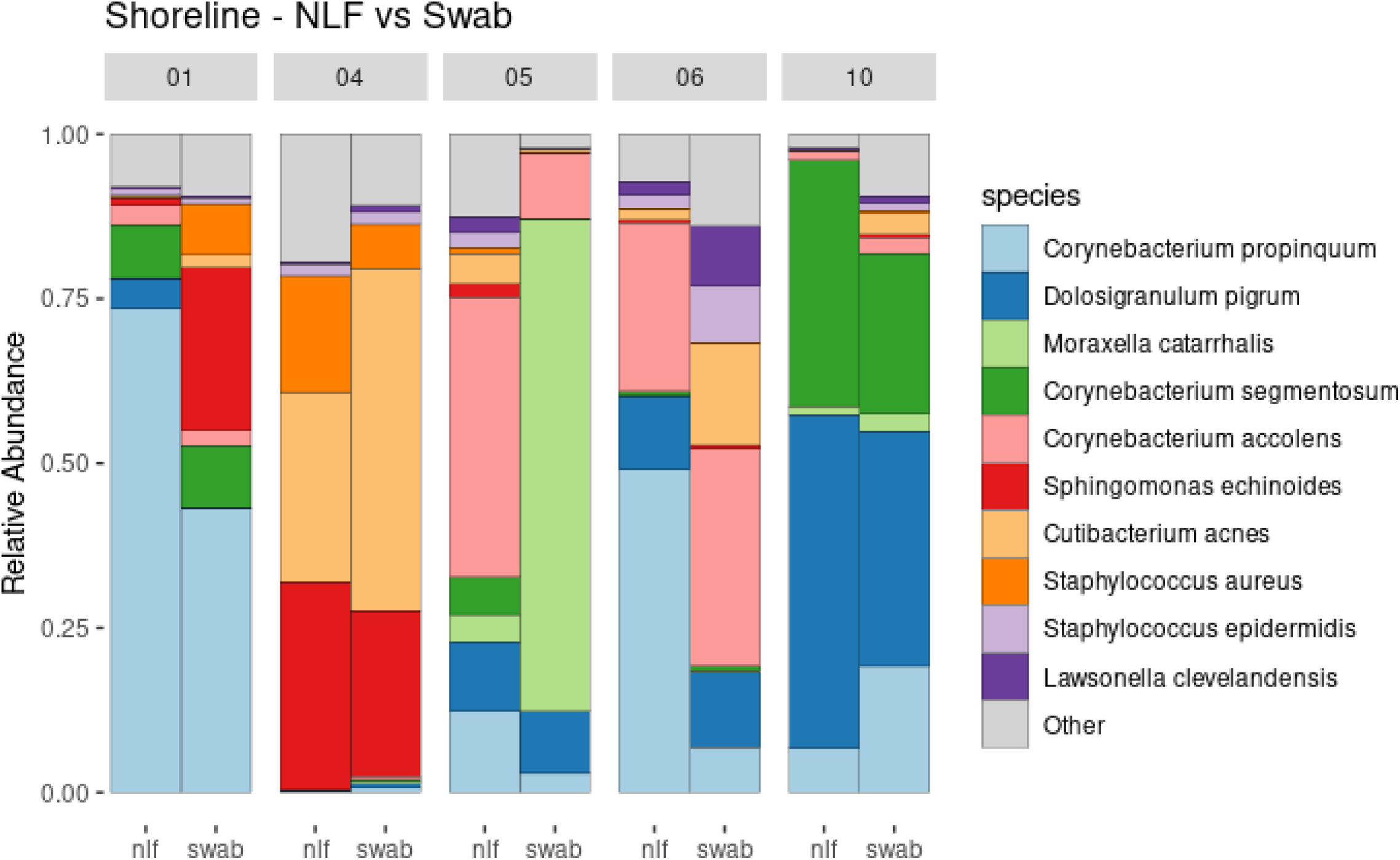

**Supplementary Fig 5.**
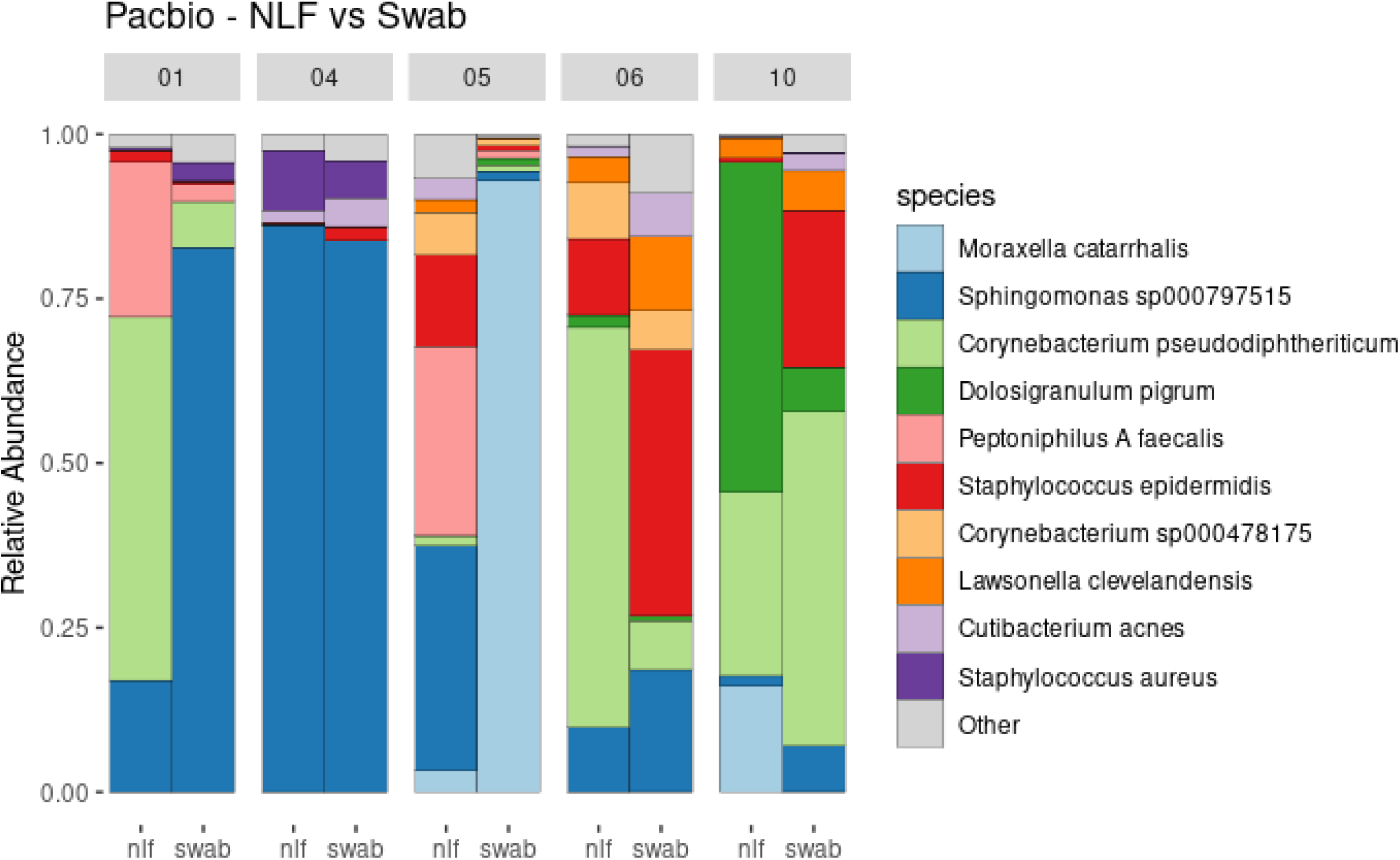

**Supplementary Fig 6.**
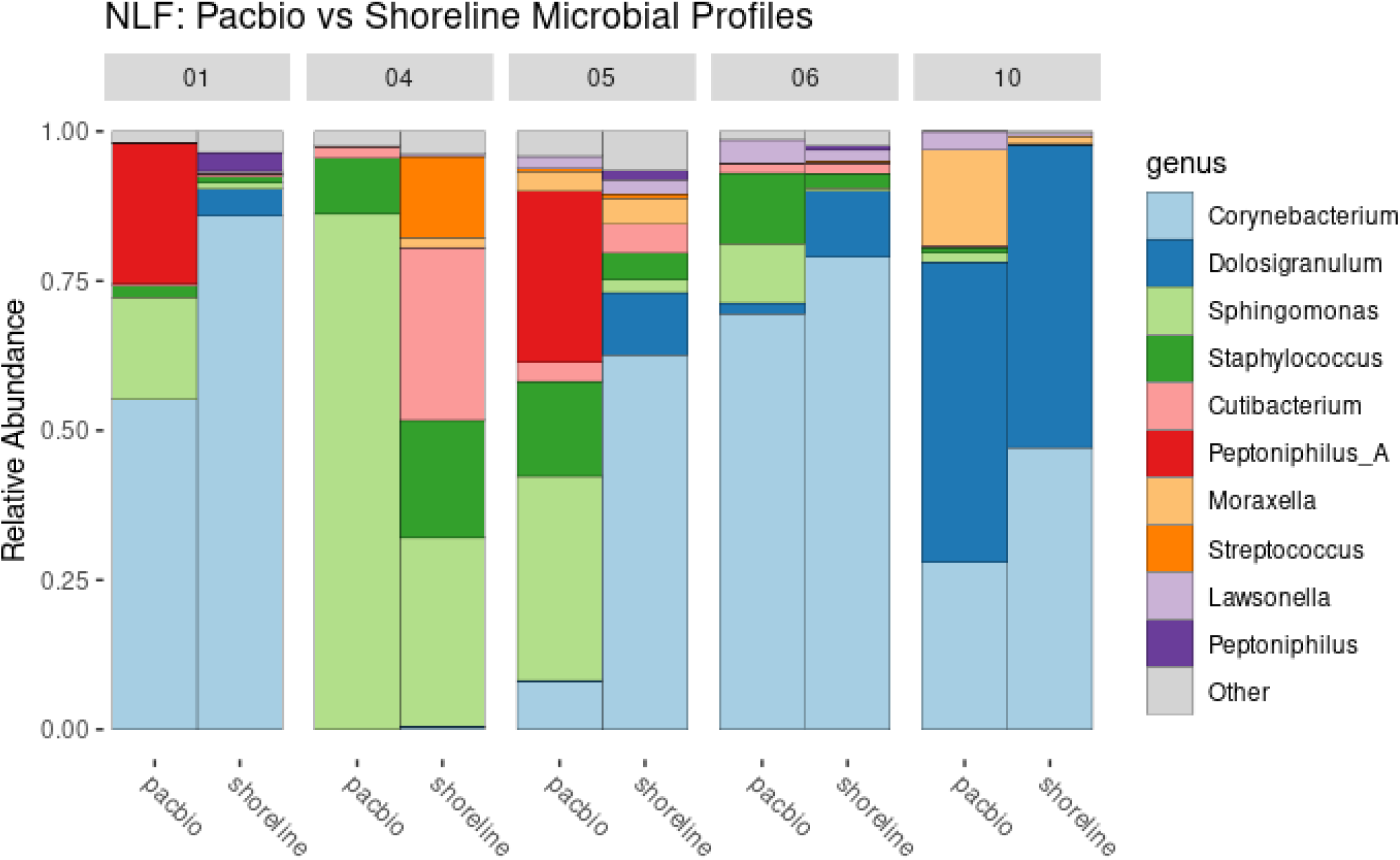

**Supplementary Fig 7.**
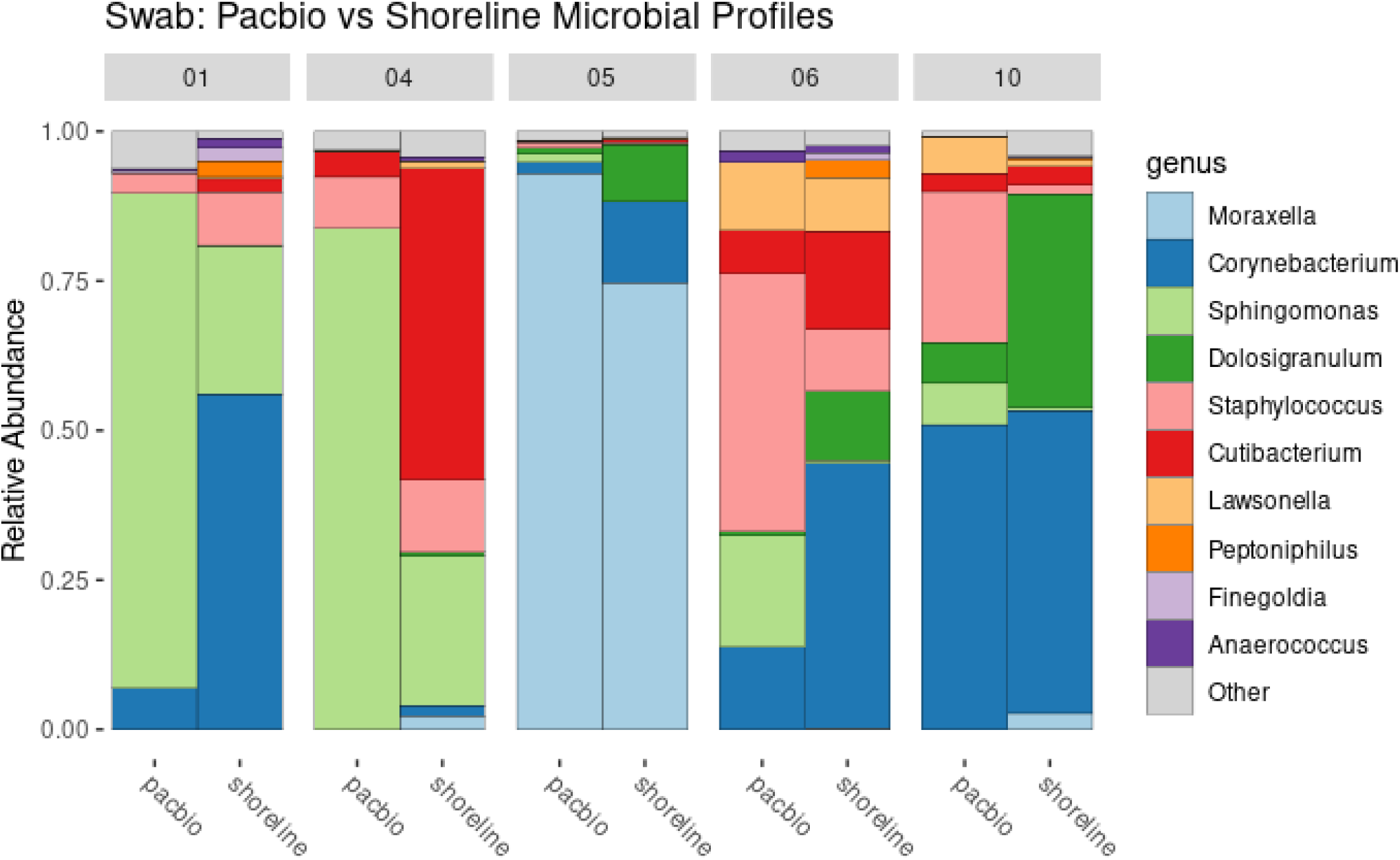

**Supplementary Fig 8.**
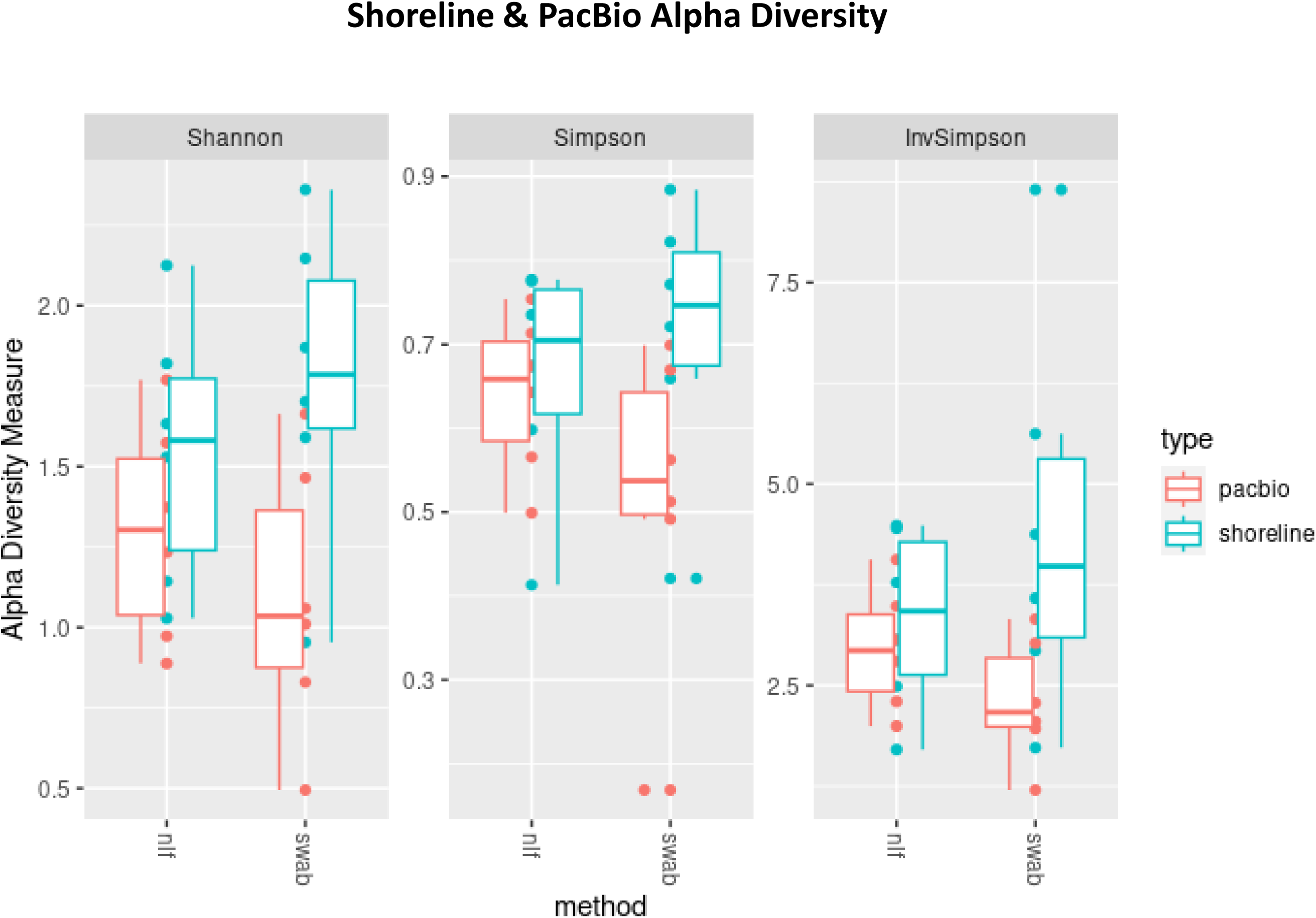

## References

1. Natalini JG, et al. The dynamic lung microbiome in health and disease. Nature Reviews Microbiology. 2023;21(4):222–35.

2. Habibi MS, et al. Neutrophilic inflammation in the respiratory mucosa predisposes to RSV infection. Science. 2020;370(6513):eaba9301.

3. Smith N, et al. Distinct systemic and mucosal immune responses during acute SARS-CoV-2 infection. Nat Immunol. 2021;22(11):1428–39.

4. Chung J, et al. Nasal lavage microbiome, but not nasal swab microbiome, correlates with sinonasal inflammation in children with cystic fibrosis. J Cyst Fibros. 2024.

5. Jochems SP, et al. Novel Analysis of Immune Cells from Nasal Microbiopsy Demonstrates Reliable, Reproducible Data for Immune Populations, and Superior Cytokine Detection Compared to Nasal Wash. PLoS ONE. 2017;12(1):e0169805.

6. Thwaites RS, et al. Nasosorption as a Minimally Invasive Sampling Procedure: Mucosal Viral Load and Inflammation in Primary RSV Bronchiolitis. J Infect Dis. 2017;215(8):1240–4.

7. Sulaiman I, et al. Functional lower airways genomic profiling of the microbiome to capture active microbial metabolism. Eur Respir J. 2021;58(1).

8. Salter SJ, et al. Reagent and laboratory contamination can critically impact sequence-based microbiome analyses. BMC Biol. 2014;12:87.

9. Erb-Downward JR, et al. Critical Relevance of Stochastic Effects on Low-Bacterial-Biomass 16S rRNA Gene Analysis. mBio. 2020;11(3).

10. Free T. Long-read sequencing for the metagenomic analysis of microbiomes. Biotechniques. 2023;74(4):153–5.

11. Gehrig JL, et al. Finding the right fit: evaluation of short-read and long-read sequencing approaches to maximize the utility of clinical microbiome data. Microb Genom. 2022;8(3).

12. Curry KD, et al. Emu: species-level microbial community profiling of full-length 16S rRNA Oxford Nanopore sequencing data. Nat Methods. 2022;19(7):845–53.

13. Albertsen M. Long-read metagenomics paves the way toward a complete microbial tree of life. Nat Methods. 2023;20(1):30–1.

14. Cuscó A, et al. Long-read metagenomics retrieves complete single-contig bacterial genomes from canine feces. BMC Genomics. 2021;22(1):330.

15. Costea PI, et al. Towards standards for human fecal sample processing in metagenomic studies. Nat Biotechnol. 2017;35(11):1069–76.

16. Sinha R, et al. Assessment of variation in microbial community amplicon sequencing by the Microbiome Quality Control (MBQC) project consortium. Nat Biotechnol. 2017;35(11):1077–86.

17. Claassen-Weitz S, et al. Optimizing 16S rRNA gene profile analysis from low biomass nasopharyngeal and induced sputum specimens. BMC Microbiol. 2020;20(1):113.

18. Pérez-Brocal V, et al. Optimized DNA extraction and purification method for characterization of bacterial and fungal communities in lung tissue samples. Scientific Reports. 2020;10(1):17377.

19. Saladié M, et al. Microbiomic Analysis on Low Abundant Respiratory Biomass Samples; Improved Recovery of Microbial DNA From Bronchoalveolar Lavage Fluid. Frontiers in microbiology. 2020;11.

20. Carney SM, et al. Methods in Lung Microbiome Research. Am J Respir Cell Mol Biol. 2020;62(3):283–99.

21. Willner D, et al. Comparison of DNA extraction methods for microbial community profiling with an application to pediatric bronchoalveolar lavage samples. PLoS ONE. 2012;7(4):e34605.

22. Thwaites RS, et al. Absorption of Nasal and Bronchial Fluids: Precision Sampling of the Human Respiratory Mucosa and Laboratory Processing of Samples. J Vis Exp. 2018(131):e56413.

23. Tighe S, et al. Genomic Methods and Microbiological Technologies for Profiling Novel and Extreme Environments for the Extreme Microbiome Project (XMP). J Biomol Tech. 2017;28(1):31–9.

24. Straub D, et al. Interpretations of Environmental Microbial Community Studies Are Biased by the Selected 16S rRNA (Gene) Amplicon Sequencing Pipeline. Frontiers in microbiology. 2020;11.

25. Ewels PA, et al. The nf-core framework for community-curated bioinformatics pipelines. Nat Biotechnol. 2020;38(3):276–8.

26. Callahan BJ, et al. DADA2: High-resolution sample inference from Illumina amplicon data. Nat Methods. 2016;13(7):581–3.

27. R Core Team. R: A language and environment for statistical computing. Vienna, Austria: R Foundation for Statistical Computing; 2021.

28. Wickham H, et al. Welcome to the Tidyverse. Journal of open source software. 2019;4(43):1686.

29. Wickham H. ggplot2: Elegant Graphics for Data Analysis. New York: Springer-Verlag; 2016.

30. Rohart F, et al. mixOmics: An R package for ‘omics feature selection and multiple data integration. PLoS Comput Biol. 2017;13(11):e1005752.

31. Davis NM, et al. Simple statistical identification and removal of contaminant sequences in marker-gene and metagenomics data. Microbiome. 2018;6(1):226.

32. Cullen JT, et al. Optimisation of a bead-beating procedure for simultaneous extraction of bacterial and fungal DNA from pig faeces and liquid feed for 16S and ITS2 rDNA amplicon sequencing. Animal - Open Space. 2022;1(1):100012.

33. Zhang B, et al. Impact of Bead-Beating Intensity on the Genus- and Species-Level Characterization of the Gut Microbiome Using Amplicon and Complete 16S rRNA Gene Sequencing. Frontiers in Cellular and Infection Microbiology. 2021;11.

34. Vesty A, et al. Evaluating the Impact of DNA Extraction Method on the Representation of Human Oral Bacterial and Fungal Communities. PLoS ONE. 2017;12(1):e0169877.

35. Shilts MH, et al. Evaluation of the upper airway microbiome and immune response with nasal epithelial lining fluid absorption and nasal washes. Scientific Reports. 2020;10(1):20618.

