## Supplementary Information for "Optimisation of DNA extraction from nasal lining fluid to assess the nasal microbiome using third-generation sequencing"

### Supplementary Methods

#### Illumina 16s rRNA sequencing

For short-read 16s rRNA amplicon-based microbial profiling, extracted DNA was amplified and sequenced by the Australian Genome Research Facility. Briefly, forward (341F: CCTAYGGGRBGCASCAG) and reverse (806R: GGACTACNNGGGTATCTAAT) primers were used to amplify the V3-4V hypervariable region of the 16s rRNA gene using conditions outlined in Supplementary Table 1. Thermocycling was completed with an Applied Biosystem 384 Veriti and using Platinum SuperFi II master mix (Invitrogen, Australia) for the primary PCR. The first stage PCR was cleaned using magnetic beads, and samples were visualised on 2% Sybr Egel (Thermo-Fisher). A secondary PCR to index the amplicons was performed with the same polymerase master mix. The resulting amplicons were cleaned again using magnetic beads, quantified by fluorometry (Promega Quantifluor) and normalised using equimolar pooling. The pooled samples were cleaned a final time using magnetic beads to concentrate the pool and then measured using a High-Sensitivity D1000 Tape on an Agilent 2200 TapeStation. The pool was diluted to 5nM and molarity was confirmed again using a Qubit High Sensitivity dsDNA assay (ThermoFisher). This was followed by sequencing on an Illumina MiSeq (San Diego, CA, USA) with a V3, 600 cycle kit (2 x 300 base pairs paired end)

| Target | Cycle | Initial | Disassociate | Anneal | Extension | Finish |
| --- | --- | --- | --- | --- | --- | --- |
| 16S: V3 - V4 | 30 | 98C for 30s | 98C for 10s | 60C for 10s | 72C for 30s | 72C for 5m |

Supplementary Table 1

### Supplementary Figures

**Supplementary Figure 1:** Quality of DNA extracted using Zymo, Qiagen, and PEG methods. **(A)** 260/280 and **(B)** 260/230 ratios of DNA extracted from the mock community standard. **(C)** 260/280 and **(D)** 260/230 ratios of DNA extracted from nasal lining fluid. * p<0.05; ** p<0.01; *** p<0.001; **** p<0.0001. Dotted line represents expected values. Differences between groups assessed using one-way ANOVA with Tukey’s test for multiple comparisons.

#### **Supplementary Figure 2:** Control samples cluster separately from true samples following 16s rRNA sequencing. **(A)** Library size (sum of reads) for control samples clearly separates from true samples. **(B)** Control samples cluster together, separately from true samples using PCoA with Bray-Curtis distance.

**Supplementary Figure 3:** Relative abundance of DNA extracted from a mock community standard. Relative abundance of genera present in **(A)** individual samples summarised in Figure 2A, **(B)** summarised by lysis method, and **(C)** summarised by extraction kit/method**.**

**Supplementary Figure 4:** Relative abundance of the ten most abundant species present in NLF and nasal swabs sequenced using Shoreline 16s rRNA sequencing.

**Supplementary Figure 5:** Relative abundance of the ten most abundant species present in NLF and nasal swabs sequenced using PacBio 16s rRNA sequencing.

**Supplementary Figure 6:** Relative abundance of the ten most abundant species present in NLF sequenced using Shoreline and PacBio 16s rRNA sequencing.

**Supplementary Figure 7:** Relative abundance of the ten most abundant species present in nasal swabs sequenced using Shoreline and PacBio 16s rRNA sequencing.

**Supplementary Figure 8:** Alpha diversity of NLF and nasal swabs sequenced using Shoreline and PacBio 16s rRNA sequencing.
